# Interspecific competition can drive the loss of conjugative plasmids from a focal species in a microbial community

**DOI:** 10.1101/2022.06.06.494957

**Authors:** David Walker-Sünderhauf, Uli Klümper, William H Gaze, Edze R Westra, Stineke van Houte

## Abstract

Plasmids are key disseminators of antimicrobial resistance genes and virulence factors, and it is therefore critical to predict and reduce plasmid spread within microbial communities. The cost of plasmid carriage is a key metric that can be used to predict plasmids’ ecological fate, and it is unclear whether plasmid costs are affected by growth partners in a microbial community. We carried out competition experiments and tracked plasmid maintenance using a synthetic and stable 5-species community and a broad host-range plasmid as a model. We report that both the cost of plasmid carriage and its long-term maintenance in a focal strain depended on the presence of competitors, and that these interactions were species-specific. Addition of growth partners increased the plasmid cost to a focal strain, and accordingly plasmid loss from the focal species occurred over a shorter time frame in these species combinations. We propose that the destabilising effect of interspecific competition on plasmid maintenance may be leveraged in clinical and natural environments to cure plasmids from focal strains.

## Introduction

Plasmids are important mobile genetic elements that facilitate horizontal gene transfer and are crucial components of microbial ecosystems. They shape microbial evolution [1,2] and are of profound clinical relevance as disseminators of antimicrobial resistance (AMR) genes [3] and virulence factors [4,5]. Many plasmids, particularly those with a broad host range, have the potential to transfer between highly diverse bacterial species [6] and mobilise resistance genes from environmental strains into clinically relevant pathogens. Hence, being able to predict and manipulate the spread of plasmids and of the genes they carry is critical to limit the spread of AMR.

A key determinant of plasmid spread and maintenance in bacterial populations and communities is the fitness effect that plasmids impose on their bacterial hosts (reviewed in [2]). Costs can arise at different steps of the plasmid lifecycle and can result, amongst others, from the expression of genes carried on the plasmid and their interference with host processes (reviewed in [7]). As a consequence, the cost of plasmid carriage varies not only between plasmids [8] but also between hosts [9]. Moreover, these costs are strongly dependent on the environment; plasmids that are costly in the absence of antibiotics or heavy metals can in turn become highly beneficial in their presence if they encode resistance genes [8].

Theory and data suggest that costly plasmids can be lost readily from bacterial populations or communities due to purifying selection, unless conjugation rates - either within or between species - are sufficiently high to support their maintenance [10–12]. For example, bacteria that lose a plasmid when cultured on their own may still associate with this plasmid when co-cultured with another species, due to high rates of interspecific plasmid transfer [10]. Hence, even bacteria that are unable to maintain plasmids in monoculture may experience increased plasmid persistence in a microbial community.

Recent work has shown that fitness effects of chromosomal mutations in bacteria can depend on the microbial community context [13,14]. We therefore hypothesised that the maintenance of plasmids in focal species may similarly be affected by the microbial community context through amplification or amelioration of the costs of carrying the plasmid. To test this hypothesis, we measured how fitness costs and maintenance of a broad host-range plasmid, pKJK5, in compost isolate *Variovorax* sp. are altered depending on the presence of competitor species in a synthetic 5-species community of soil bacterial isolates. These are *Pseudomonas* sp., *Stenotrophomonas* sp., *Achromobacter* sp., and *Ochrobactrum* sp., which form a stable microbial community together with *Variovorax* sp. [15,16]. After finding that the cost of carrying pKJK5 to *Variovorax* was increased in the presence of certain growth partners, we measured plasmid maintenance in a series of synthetic communities of varying complexity. The pattern of pKJK5 maintenance was predictable based on the costs of plasmid carriage; *Variovorax* lost pKJK5 more rapidly when embedded in the community. Finally, we generalised community-dependent plasmid loss by measuring pKJK5 maintenance in each constituent of the microbial community and found additional examples where pKJK5 maintenance in a constituent species depended on the microbial community context.

## Results

### Plasmid cost to *Variovorax* increased in presence of growth partners

We hypothesised that the cost of carrying the broad-host-range, IncP-1ε plasmid pKJK5 in *Variovorax* sp. depends on the microbial community context. To test this, we measured the fitness costs of carrying the plasmid by competing pKJK5-bearing *Variovorax* that was chromosomally tagged with Gentamicin resistance gene *aacC1* (V(Gm^R^)) with pKJK5-free *Variovorax* that was chromosomally tagged with Chloramphenicol resistance gene *catR* (V(Cm^R^)). These clones were competed either in isolation or in the presence of each of the four different growth partners (*Pseudomonas* sp., *Stenotrophomonas* sp., *Achromobacter* sp., or *Ochrobactrum* sp.) in 5 biological replicates. To enable visualisation of plasmid transfer, we engineered pKJK5::*gfp*^PL^, which encodes green fluorescent protein (GFP). Furthermore, we inserted payload (PL) genes into the plasmid to increase the baseline cost of plasmid carriage to the host (Figure S1b). As a control, we also competed pKJK5-free V(Gm^R^) with pKJK5-free V(Cm^R^) in each of these contexts.

These direct competition experiments revealed that pKJK5::*gfp*^PL^ carriage was associated with a fitness cost to *Variovorax* in monoculture (Figure 1). The plasmid’s selection coefficient was not altered in the presence of *Achromobacter* or *Ochrobactrum*. However, the presence of either *Pseudomonas* or *Stenotrophomonas* significantly decreased the selection coefficient of pKJK5::*gfp*^PL^ (p<0.001 and p<0.05 respectively; Gaussian GLM, F=20.71, d.f.=9&140, adjusted R^2^=0.54). This demonstrates that carrying pKJK5::*gfp*^PL^ was more costly to *Variovorax* in the presence of *Pseudomonas* or *Stenotrophomonas* compared to *Achromobacter* or *Ochrobactrum* or in monoculture.

**Figure 1:**
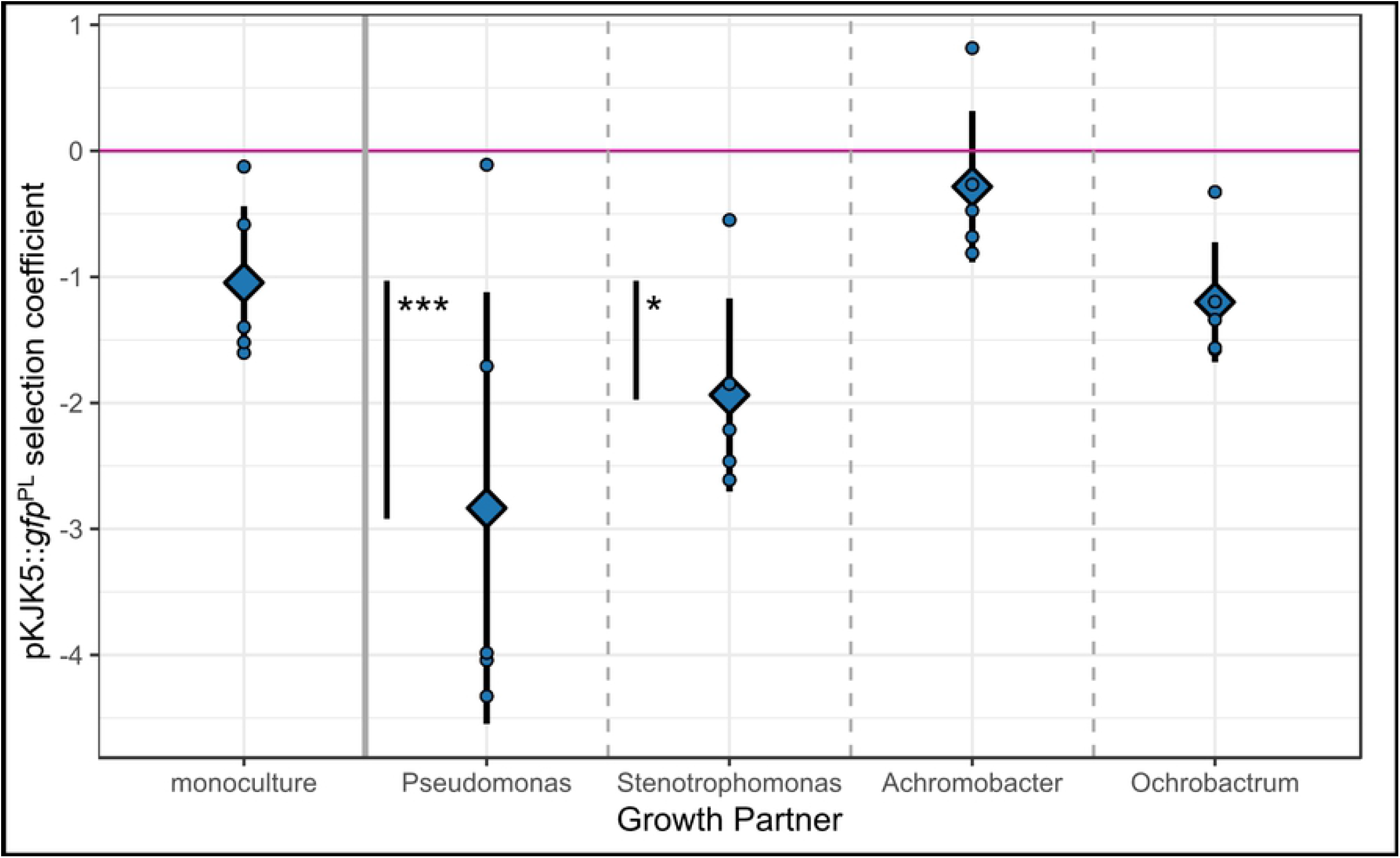
Addition of growth partners alters pKJK5::*gfp*^PL^’s selection coefficient. Mean ± standard deviation (including individual datapoints) of pKJK5::*gfp*^PL^ selection coefficient to *Variovorax* in monoculture and with different growth partners after 5 days of co-culture. Values >0 indicate a fitness benefit and values <0 indicate a fitness cost of carrying pKJK5::*gfp*^PL^ in each context. ***p<0.001, *p<0.05 as calculated by Tukey’s Honest Significance Difference test (HSD) after fitting a Gaussian generalised linear model (GLM); F=20.71; d.f.=9 & 140; adjusted R^2^=0.54; N=5.

Furthermore, analysis of GFP expression of *Variovorax* colonies on various selective plates revealed that the frequencies of plasmid conjugation were very low under our experimental conditions, with only 8 out of 983 colonies of the initially pKJK5-free *Variovorax* strain becoming GFP positive (~0.8% transconjugants/recipients; Table S1).

### Plasmid costs were linked to maintenance

Based on these results, we hypothesised that the increased fitness cost of pKJK5::*gfp*^PL^ to *Variovorax* in the presence of *Pseudomonas* and *Stenotrophomonas* would also lead to decreased plasmid maintenance. To test this, we generated 16 different synthetic microbial communities with 5 replicates each, composed of all possible combinations of one-, two-, three-, four-, and five species, consisting of *Variovorax* carrying pKJK5::*gfp*^PL^ and pKJK5-free *Pseudomonas, Stenotrophomonas, Achromobacter*, and/or *Ochrobactrum* (Figure S2). We measured plasmid maintenance after 5 days of co-culture. In monoculture, nearly all *Variovorax* clones in the population retained pKJK5::*gfp*^PL^ (Figure 2). In contrast, plasmid maintenance in *Variovorax* was significantly decreased in seven of the 16 synthetic communities (p<0.05; binomial GLM, F=0.46, d.f.=17&62, pseudo R^2^=0.85, additional details in Figure 2 and Table S2). Interestingly, these corresponded to all those communities that contained *Stenotrophomonas*, with the sole exception of the full 5-species community where the reduction in *Variovorax* plasmid maintenance was not significant (p=0.65).

**Figure 2:**
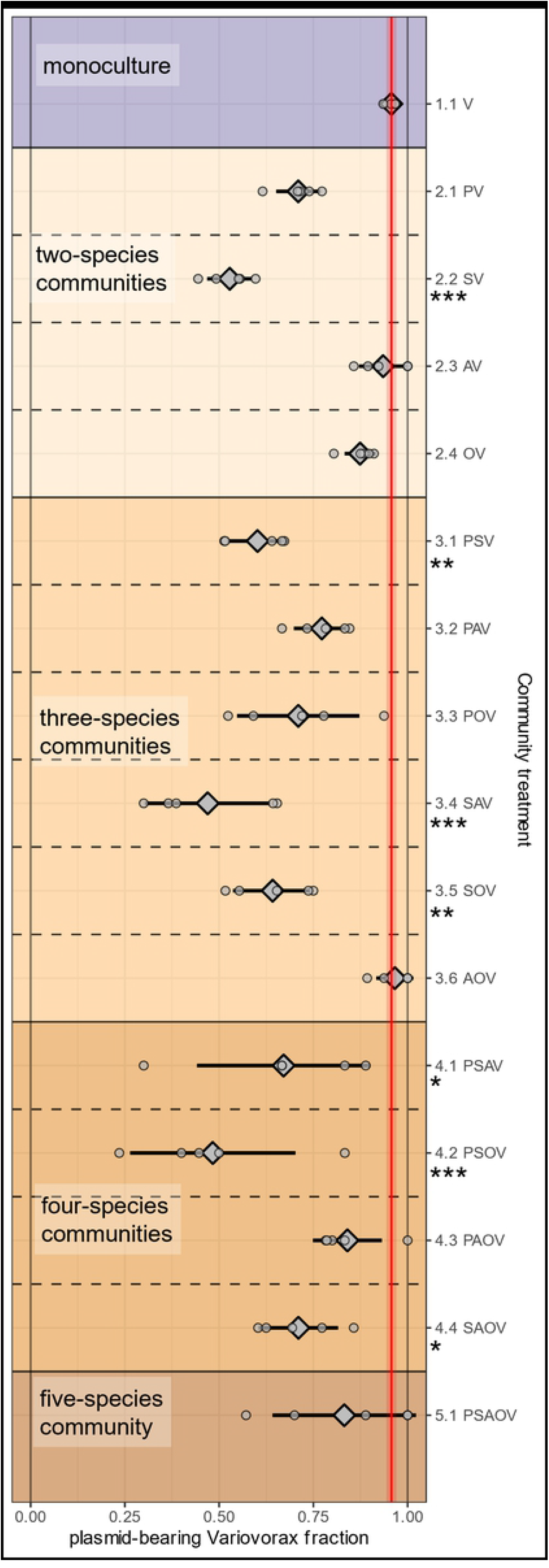
Different combinations of growth partners elicit different *Variovorax* plasmid loss effects. Mean ± standard deviation (including individual datapoints) of GFP+ *Variovorax* (V) fraction as a proxy of plasmid-bearing *Variovorax* in the presence of various growth partners (*Pseudomonas* (P), *Stenotrophomonas* (S), *Achromobacter* (A), *Ochrobactrum* (O)) after 5 days of co-culture. For comparison, the vertical red line and shaded areas indicate the mean ± standard deviation of the monoculture treatment. Stars indicate treatments with significantly lower GFP+ fraction than in monoculture. See Table S2 for values of these summary statistics. *p<0.05, **p<0.01, ***p<0.001 as calculated by Tukey’s HSD after fitting a binomial GLM; F=0.46; d.f.=17&62; pseudo R^2^=0.85; N=5.

We explored how the proportion of each constituent species correlated with the proportion of pKJK5-bearing *Variovorax* in each replicate of the 16 communities by fitting separate linear models (LMs) to each subset of treatments, excluding those which lacked the species being investigated (Figure 3). The proportions of *Stenotrophomonas* (p = 0.042), *Achromobacter* (p = 0.00013), and *Variovorax* (p = 0.025) significantly associated with the plasmid-maintaining fraction of *Variovorax* within the same communities, the same was not true for the proportions of *Pseudomonas* (p = 0.90) or *Ochrobactrum* (p = 0.095). Upon closer investigation, *Variovorax* community proportion was associated with plasmid maintenance only due to the monoculture treatment, without which no correlation existed (p = 0.98). Additionally, we found that several species proportions, such as *Stenotrophomonas* and *Achromobacter*, co-correlated (Figure S3). Therefore, this simplistic analysis of species proportion and plasmid maintenance had several shortcomings including an effect driven by a single monoculture treatment, correlation between individual variables, removal of different subsets of data when excluding absent species, and multiple statistical testing. To address all these issues, we instead fitted a single statistical model to all available data, which we used above to analyse the various treatments in Figure 2. This also revealed that only *Stenotrophomonas* and *Achromobacter* proportion had statistically significant effects on *Variovorax* plasmid maintenance (Table S3). There was a clear negative association between the proportion of *Stenotrophomonas* and the plasmid-bearing *Variovorax* fraction. Interestingly, *Achromobacter* had the opposite effect: its presence was associated with higher plasmid maintenance. While part of these differentiating effects may be explained by the negative correlation between *Stenotrophomonas* and *Achromobacter* proportions (Figure S3), the different impacts these species have on plasmid selection coefficient of pKJK5::*gfp*^PL^ to *Variovorax* in coculture (Figure 1) suggest that a change in plasmid fitness costs plays a role, too.

**Figure 3:**
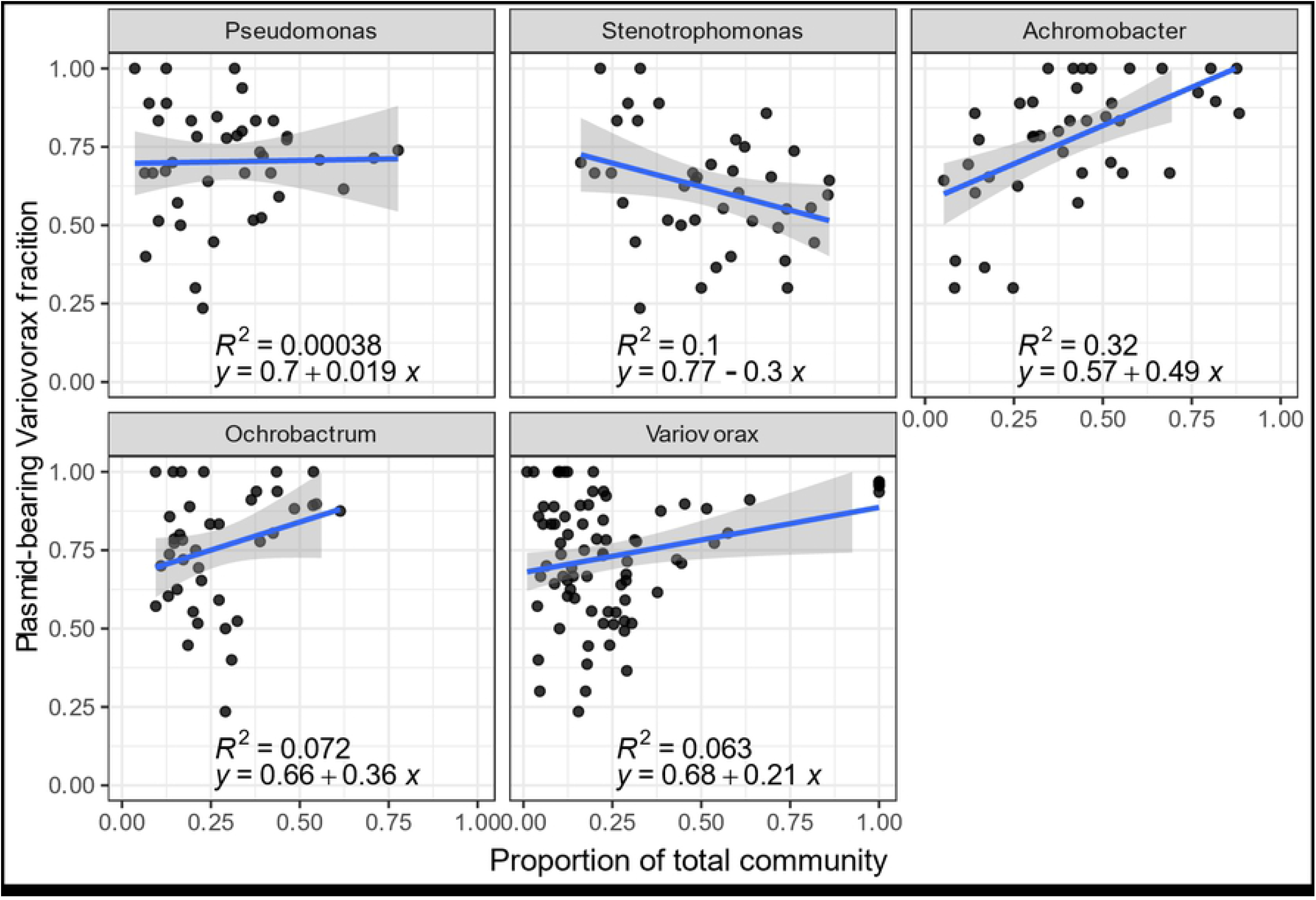
The relationship of community composition and *Variovorax* plasmid maintenance. Community composition of all samples of various communities as in Figure 2 plotted against the GFP+ fraction of *Variovorax* colonies as a proxy of pKJK5-bearing *Variovorax*. Community composition is broken up into individual panels describing *Pseudomonas, Stenotrophomonas, Achromobacter, Ochrobactrum*, and *Variovorax* proportions of the whole community respectively. Counts of 0 for each community constituent were removed for each panel. Blue lines and shaded areas indicate fitted LMs with equations and R^2^ displayed in each panel. Note these relationships are for illustrative purposes only, and not used to determine statistical significance to avoid multiple testing. Of these five metrics, only *Stenotrophomonas* and *Achromobacter* proportion constitute significant terms of the statistical model fitted to the data; see Table S3.

To formally address the hypothesis that the differences in plasmid maintenance across the synthetic communities were caused by differences in the cost of plasmid carriage to *Variovorax*, we performed a correlational analysis between the selection coefficient of pKJK5::*gfp*^PL^ to *Variovorax* (Figure 1) and maintenance of this plasmid (Figure 2). Treatments in which the *Variovorax* plasmid selection coefficient was higher also displayed higher levels of *Variovorax* plasmid maintenance (Figure 4; p=0.023, mixed-effects LM, F=5.26, 25 observations, 2x 5 random-effect groups, conditional R^2^=0.18). The monoculture treatment and community treatments consisting of either *Achromobacter* or *Ochrobactrum* as a growth partner all clustered in the top-right quadrant, representing treatments where plasmid cost was low and plasmid maintenance high. In contrast, community treatments that contained either *Pseudomonas* or *Stenotrophomonas* as a growth partner were associated with low plasmid maintenance and high costs of plasmid carriage for *Variovorax*.

**Figure 4:**
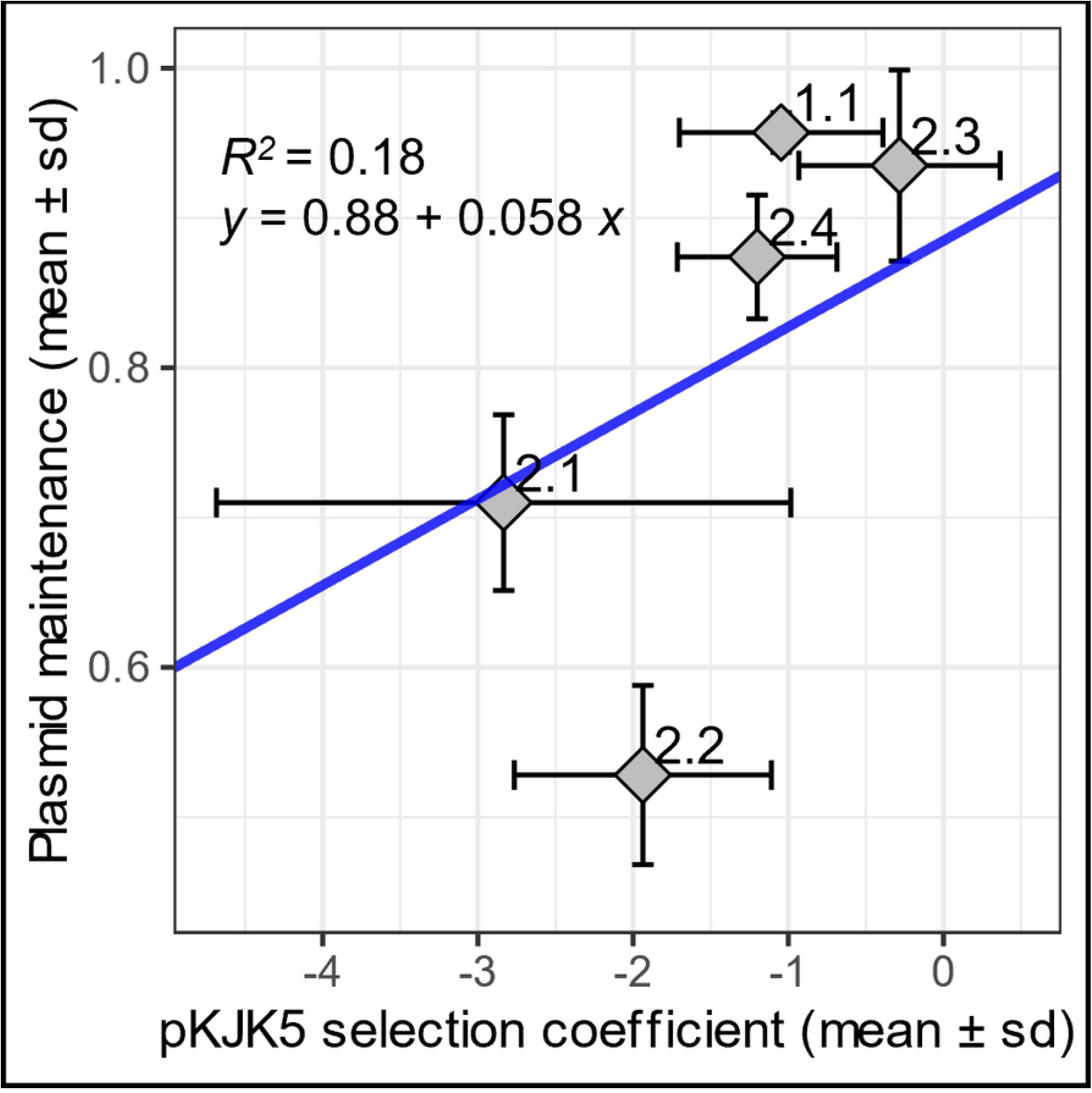
*Variovorax* pKJK5::*qfp*^PL^ selection coefficienct correlates with plasmid maintenance. Mean ± standard deviation of pKJK5::*gfp*^PL^ selection coefficient as in Figure 1 and pKJK5-bearing *Variovorax* fraction in corresponding treatments as in Figure 2. Treatment designations indicate growth partners: 1.1 monoculture, 2.1 *Pseudomonas*, 2.2 *Stenotrophomonas*, 2.3 *Achromobacter*, 2.4 *Ochrobactrum*. Blue line indicates fitted mixed-effects linear model with equation and R^2^ as displayed. F=5.26, 25 observations with 2x 5 random-effect groups. Selection coefficient is a significant determinant of plasmid-bearing *Variovorax* fraction, p=0.023.

### pKJK5 maintenance was community-dependent in multiple species

Based on these results, we sought to explore whether the observed effects were unique to *Variovorax* and pKJK5::*gfp*^PL^ or if they could be observed in other host-plasmid pairs. To exclude the possibility that plasmid maintenance in the presence of growth partners may have been influenced by the additional payload genes inserted into pKJK5::*gfp*^PL^ (Figure S1b) and to expand the analysis to a plasmid with smaller predicted fitness cost, we used plasmid *pKJK5::gfp* which lacks these payload genes (Figure S1c [6]). We therefore carried out plasmid maintenance experiments in which all species carried the plasmid (*Pseudomonas, Stenotrophomonas, Achromobacter, Ochrobactrum, Variovorax*), either with the presence or absence of additional payload genes (pKJK5::*gfp*^PL^ and pKJK5::*gfp*, respectively). The frequencies of pKJK5::*gfp*^PL^ and of pKJK5::*gfp* carriage were measured over 17 days in each host cultured in isolation or as part of a stable microbial community consisting of all five species bearing the plasmid. For both pKJK5 variants, maintenance of the plasmid strongly depended on host identity; *Ochrobactrum* retained pKJK5::*gfp* and pKJK5::*gfp*^PL^ to high levels, while *Pseudomonas* and *Achromobacter* lost these plasmids during the course of the experiment (Figure 5). Notably, these differences in plasmid maintenance were independent of the community context in which the hosts were cultured.

**Figure 5:**
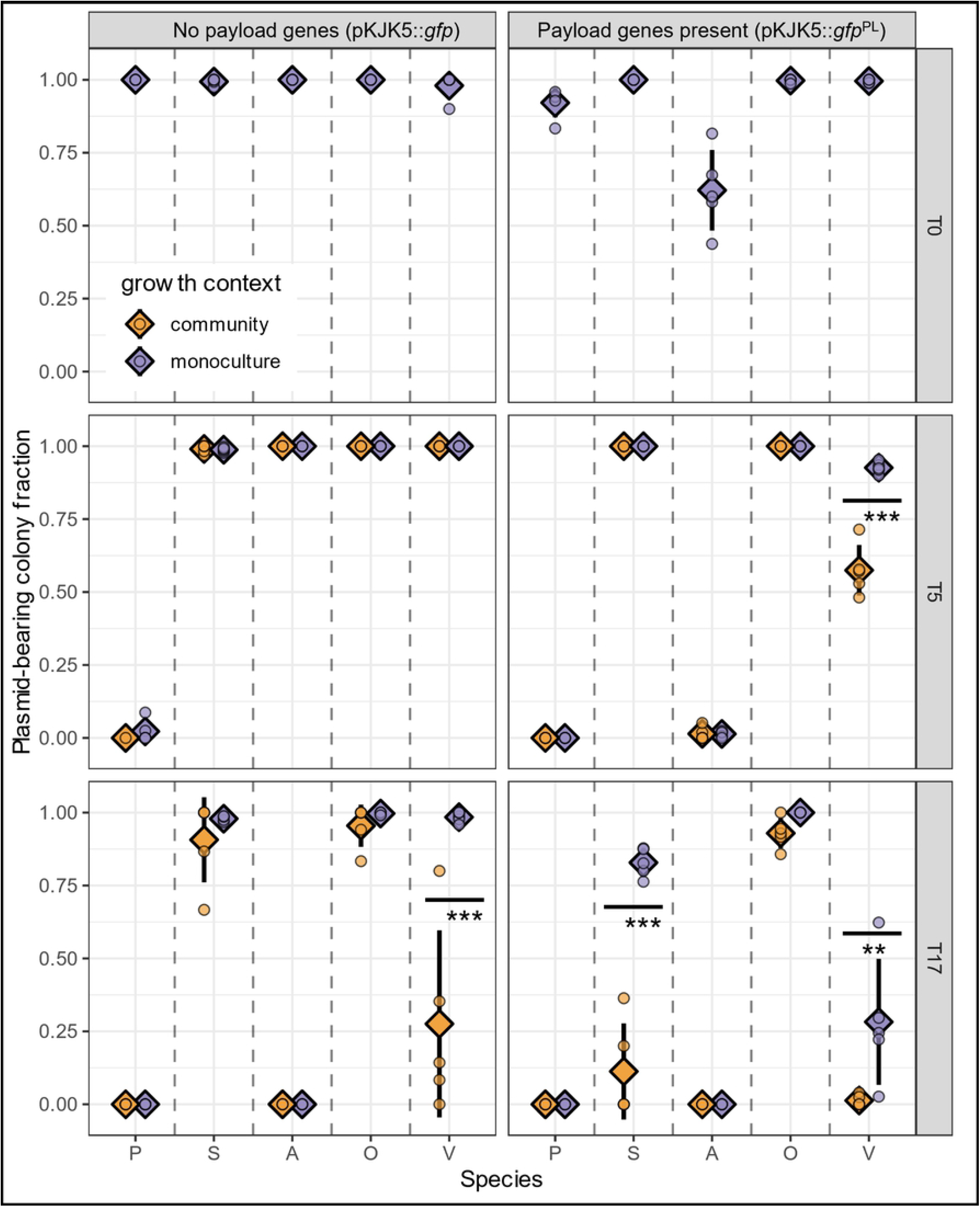
*Variovorax* and *Stenotrophomonas* pKJK5 maintenance is dependent on community context. Mean ± standard deviation (including individual datapoints) of GFP+ fraction of colonies as a proxy of pKJK5::*gfp* (left) or pKJK5::*gfp*^PL^ (right) maintenance. Data is split into species *Pseudomonas* (P), *Stenotrophomonas* (S)*, Achromobacter* (A), *Ochrobactrum* (O), and *Variovorax* (V) after 5 and 17 days of growth in monoculture or in a community context. T0 indicates pKJK5-maintaining proportion of strains used to start the experiment. **p<0.01, ***p<0.001 as calculated by Tukey’s HSD after fitting binomial GLMs; T5: F=1.49; df=13 & 73; pseudo R2= 0.99; T17: F=0.64; df=19 & 74; pseudo R2=0.98; N=4-5. No other treatment combinations are significantly different from each other.

As expected, *Variovorax* plasmid maintenance was dependent on the community context it is cultured in. In monoculture, 93±2% of colonies retained pKJK5::*gfp*^PL^ after 5 days. In contrast, significantly fewer (57±9%) *Variovorax* clones within the community retained pKJK5::*gfp*^PL^ (p<0.001; binomial GLM; F=1.49; df=13 & 73; pseudo R^2^= 0.99). This effect remained evident after 17 days of culture (28±22% *Variovorax* plasmid maintenance in monoculture, 1.3±1.9% maintenance in community context). Similar dynamics could be observed for *Variovorax* carrying *pKJK5::gfp* after 17 days, where 98±2% of colonies retained pKJK5::*gfp* in monoculture compared to 28±32% of colonies in a community context. As significant pKJK5::*cas*^PL^ plasmid loss from *Variovorax* had not been observed when growth partners did not carry the plasmid (Figure 2; treatment 5.1), these results also highlight how context-dependent plasmid maintenance was: whether or not growth partners also carried a plasmid had an impact on community structure, which in turn affected pKJK5::*cas*^PL^ maintenance outcome in *Variovorax*.

Plasmid maintenance in *Stenotrophomonas* was very high for both pKJK5 variants after 5 days, independent of community context. However, after 17 days, *Stenotrophomonas* pKJK5::*gfp*^PL^ maintenance remained significantly higher in monoculture (83±5%) than in a community context (11±16%; p<0.001; binomial GLM; F=0.64; df=19 & 74; pseudo R^2^=0.98). While no statistical difference of *Stenotrophomonas pKJK5::gfp* maintenance was observed, average maintenance of *pKJK5::gfp* was slightly lower in community context after 17 days of growth (91±15% maintenance in community vs. 98±1% maintenance in monoculture).

Collectively, the baseline cost of plasmid carriage led to quantitatively different but qualitatively similar plasmid maintenance dynamics in all host species. pKJK5::*gfp*^PL^’s payload genes *cas9* and non-targeting *sgRNA* (Figure S1b) are known to carry a constitutive cost to the bacterial host [17–19]. The smaller payload of pKJK5::*gfp* slowed plasmid loss dynamics in comparison to costlier pKJK5::*gfp*^PL^. Furthermore, the presence of a microbial community accelerated loss of plasmid pKJK5::*gfp*^PL^ in multiple species of this synthetic community.

## Discussion

This work aimed to understand how costs and maintenance of a conjugative plasmid can be determined by the microbial community context of its host by using a model community composed of five different species of soil bacteria. These species form a locally maladapted community in which the most common form of community interactions is resource competition. No bacterial warfare in form of direct growth inhibition or killing of growth partners takes place in this system, and all community members can grow to an equilibrium concentration when invading from rare [15,16]. Under culture conditions which favour long-term species coexistence, little to no conjugation took place (Table S1).

Using this model community, we found that the fitness cost of plasmid carriage can change as a result of interspecific competition. We found that embedding focal strain *Variovorax* sp. in a community led to increased costs of pKJK5::*gfp*^PL^ plasmid carriage and more rapid loss of this plasmid. The mechanism of plasmid loss in our system is unknown; but both higher intrinsic plasmid loss rates and higher relative growth of plasmid-free segregants would lead to the same observed outcome of population-level plasmid loss, and both can be driven by increased cost of plasmid carriage. This increased cost may be a result of resource limitation during growth with competitors. Generally, plasmid maintenance is dependent not only on fitness costs and benefits to its host, but also the plasmid transfer and loss rate [20,21]. Of these, plasmid transfer can also depend on the community context: In monoculture, a mercury resistance plasmid was rapidly lost from its host species, but with growth partners, the plasmid was maintained in the focal strain due to reinfection by conjugation [10,22]. This phenomenon of plasmid persistence through conjugation has been observed for multiple types of plasmids, and in communities consisting of several *Escherichia coli* strains and several plasmids [21]. Ultimately, in these experiments, embedding a plasmid host in a community context led to increased plasmid maintenance as a result of increased conjugation. Overall, the fitness cost of the plasmids was not a good predictor of plasmid maintenance [10]. In contrast, in our experiments the plasmid selection coefficient was a significant determinant of plasmid maintenance outcome – therefore, we propose that a trade-off of plasmid cost and plasmid conjugative transfer may determine plasmid maintenance outcome. Which of these variables has higher importance depends on the study system: We speculate that plasmid maintenance in a focal species is increased in communities with high plasmid conjugation frequencies by providing a reservoir of plasmid-containing cells that re-infect other bacteria in the community. In turn, plasmid maintenance is decreased in communities with low conjugation frequencies, where interspecific competition can increase the cost of plasmid carriage. That said, recent work indicates that community context can also limit conjugation to focal species due to the dilution effect [23].

Overall, this study and previous work emphasise the importance of variable plasmid costs and plasmid re-infection rates when considering plasmid maintenance in a focal host species within a community context.

It is likely that other species beyond our model system maintain plasmids in a community-dependent manner. Provided we have sufficient knowledge about community interactions and plasmid behaviour to predict the outcome of embedding a focal strain in a community context, community-dependent plasmid loss could have wider implications: In healthcare, bacteria might lose a plasmid containing virulence factors or antibiotic resistance genes in a new community context. For instance, a plasmid present in environmental microbiomes might become lost once the plasmid host becomes embedded in a human microbiome, as observed when a virulence plasmid became lost from *E. coli* in the human gut during the course of an infection [24], although the mechanisms behind plasmid loss were not investigated in this study. Conversely, resistance plasmids which are maintained in a community context may be lost in diagnostic monocultures. Community-level plasmid loss would especially need to be investigated if previously characterised highly permissive plasmid hosts, which are key to plasmid maintenance on a community level [25], are shown to experience community-dependent impacts on their plasmids’ fitness cost.

Further, this work could have implications on the spread of synthetic plasmids bearing payload genes through microbial communities (e.g. CRISPR-Cas antimicrobials [26]). Where payload genes increase the plasmid’s fitness cost, this may impact its maintenance in a microbial community even when the plasmid can be stably maintained in single bacterial strains. Finally, this work opens exciting research avenues for manipulation of plasmid content of focal species. For example, removal of virulence and resistance plasmids from pathogens may be achieved through addition of certain plasmid loss-inducing growth partners. This prediction needs to be tested thoroughly using additional plasmid-host pairs.

## Materials and Methods

### Strains and culture conditions

Focal strain *Variovorax* sp. (V) forms a synthetic community with bacterial compost isolates *Pseudomonas* sp. (P), *Stenotrophomonas* sp. (S), *Achromobacter* sp.(A), and *Ochrobactrum* sp.(O). *Variovorax* is a β-proteobacterium, and members of this genus are often found in microbial soil communities. The model species form a stable community *in vitro* over very long timescales (>1 year) and form visually distinct colonies on King’s B medium (KB) agar, allowing to enumerate species frequencies without the need for selective plating [15]. All communities and monocultures were incubated in 6 mL low-nutrient 1/64 Tryptone Soy Broth (TSB; diluted in water) statically at 28°C. For analysis, samples were plated onto KB agar plates at appropriate dilutions and incubated at 28°C for 2-3 days. Community composition was assessed by counting each colony phenotype, ambiguous colony phenotypes were confirmed by 16S colony PCR and Sanger sequencing (using primers 27F 5’-agagtttgatcmtggctcag-3’ and 1492R 5’-accttgttacgactt-3’). Plasmid carriage was assessed by screening GFP expression using a fluorescence lamp (NightSea lamp with RB bandpass filter).

### Engineering of pKJK5::*gfp*^PL^ conjugative plasmid

pKJK5 is a 54-kb IncP1-ε plasmid originally isolated from a manure-associated microbial soil community that carries resistance genes to Tetracycline, Trimethoprim, Aminoglycosides, and Sulfonamides within an *intI1* integron cassette [27]. It can readily be transferred into highly diverse members of soil or waste-water treatment plant communities [6,25].

Using pKJK5 as a template, conjugative plasmid pKJK5::*gfp*^PL^ was constructed by 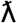-red mediated recombineering using pACBSCE and pDOC_Cas in *E. coli* DH5α with a protocol adapted from [28]: we inserted a gene cassette constitutively expressing GFP, *Streptococcus pyogenes* Cas9 (the SpyCas9 sequence described in [29] was codon-optimised to pKJK5 codon usage), and sgRNA into pKJK5’s *intI1* gene (Figure S1b). The sgRNA targets a random nucleotide sequence absent from the study system (gttttctgcctgtcgatcca). Alongside this, we used pKJK5::*gfp* [6] (Figure S1c). pKJK5::*gfp*^PL^ and pKJK5::*gfp* were delivered to P, S, A, O, and V using *E. coli MFDpir* [30] or *E. coli* K12::mCherry [6] as donors, followed by selecting GFP-expressing (GFP+) transconjugant colonies on KB agar plates containing 12 μg/mL Tetracycline only (pKJK5::*gfp*) or Tetracycline and 10 μg/mL Trimethoprim (pKJK5::*gfp*^PL^).

### *Variovorax* fitness experiment (Figure 1)

Two chromosomal tag variants of *Variovorax* were constructed using mini-Tn5-transposon vectors pBAM1-Gm [31] and derivative pBAM1-Cm which contains Chloramphenicol resistance gene *catR* [32]. These suicide transposon vectors were delivered to *Variovorax* using auxotrophic donor strain *E. coli* MFD*pir* following established protocols [32]. Successful insertion of *aacC1* and *catR* genes was confirmed by their resistant phenotype and by PCR.

Five replicate V(Gm^R^) + pKJK5::*gfp*^PL^ transconjugant colonies (generated as described above) were suspended in 1/64 TSB + 12μg/mL Tetracycline, and five replicate colonies each of plasmid-free V(Gm^R^), V(Cm^R^), P, S, A, and O were suspended in 1/64 TSB. After initial two-day incubation, cultures were washed twice with 0.9% (w/v) NaCl solution to remove all traces of antibiotics before being used to start the experiment. Communities were established by using 20 μL of each *Variovorax* strain (adjusted to OD600=0.065) mixed with 50μL of OD-adjusted P, S, A, or O. For the pKJK5-bearing treatment, V(Gm^R^)+pKJK5::*gfp*^PL^ and V(Cm^R^) competed against each other either alone or together with P, S, A, or O. In the pKJK5-free control, V(Gm^R^) and V(Cm^R^) competed against each other in the same contexts. All competitions were carried out in the absence of selection.

The communities were cultured for three days, vortexed and transferred into fresh microcosms and incubated for another two days. Then, communities were vortexed and 50 μL of a 10^-5^ dilution of each sample was plated onto KB, KB+50μg/mL Gentamicin, and KB+25μg/mL Chloramphenicol plates.

Robustness of chromosomal tags was confirmed by colony PCR of *aacC1* and *catR* of 66 random *Variovorax* colonies across treatments (using primers aacC1-fw 5’-atgttacgcagcagcaacga-3’; aacC1-rv 5’-ttaggtggcggtacttgggt-3’; cm-fw 5’-agacggcatgatgaacctga-3’; cm-rv 5’-cggtgagctggtgatatggg-3’). Colony identities of all species were assessed on each plate. The relative fitness of V(Gm^R^) and pKJK5 selection coefficient *s* for each treatment were calculated using the following equations:

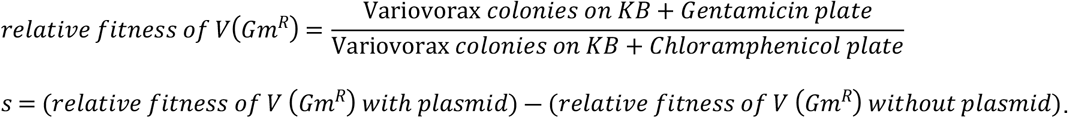

### Plasmid maintenance experiments (Figures 2–5)

To set up monocultures and communities for the plasmid maintenance experiments (Figures 2 and 5), five colonies of each community constituent were individually suspended in 1/64 TSB, supplemented with 12 μg/mL Tetracycline where the community constituent carried pKJK5::*gfp*^PL^ or pKJK5::*gfp*. Adding Tetracycline at this step ensured pKJK5 maintenance in all strains except *Achromobacter*, which displays resistance to this antibiotic even in the absence of pKJK5.

After 2 days incubation, 1 mL of these cultures was pelleted and washed twice with 0.9% NaCl to remove all traces of antibiotics. For monocultures, 20 μL of each of the five separately cultured colonies per isolate were transferred into fresh microcosms, giving rise to five biological replicate monocultures per treatment. Additionally, 15 community combinations were established by mixing 20 μL of each five replicates of P, S, A, and/or O and V, giving rise to five replicate communities per treatment (Figure S2b). All experiments were carried out in the absence of selection.

Monocultures and communities were cultured for three days, vortexed, and 100 μL of each culture transferred into a fresh microcosm and incubated for another two days. Communities were then further passaged into fresh microcosms the same way every two days until 17 total days of co-culture (Community Maintenance experiment only). To assess community composition, 50 μL of a 10^-5^ dilution of samples were plated onto KB agar plates at T0, T5, and T17 (Figure S2c).

### Statistical analyses

Data processing, data visualisation, and statistical analyses were carried out using R version 4.0.5 and RStudio version 1.4.1103 with the following packages: dplyr, tidyr, readr, ggplot2, ggpubr, lme4, GGally, and MASS.

For all analyses, other model types, link functions, and the inclusion of additional variables were tested. The final models were found to be the most robust. Model assumptions were checked and found to be upheld. For comparison of specific treatment categories, Tukey’s post-hoc test of honest significance differences was carried out. The model details are as follows:

#### Variovorax *fitness (Figure 1)*

We constructed a gaussian generalised linear model (GLM) with logit link function: Plasmid selection coefficient was modelled as a function of Treatment, Growth Partner, and their interaction. F=20.71, df=9 & 140, adjusted R^2^=0.5435.

#### Growth partner Experiment (Figure 2)

Initially, we explored the data by fitting 5 individual LMs, each describing *Variovorax* plasmid maintenance as a function of species prevalence. For each model, datapoints with a prevalence of 0 were dropped. *Pseudomonas* proportion: F=0.015; df=1&38; p=0.90; R^2^=0.00038. *Stenotrophomonas* proportion: F=4.44; df=1&38; p=0.042; R^2^=0.10. *Achromobacter* proportion: F=18.11; df=1&38; p=0.00013; R^2^=0.32. *Ochrobactrum* proportion: F=2.94; df=1&38; p=0.095; R^2^=0.072. *Variovorax* proportion: F=5.21; df=1&78; p=0.025; R^2^=0.063.

Fitting a LM between *Variovorax* proportion and plasmid maintenance without the monoculture data showed no association: F=0.0006; df=1&73; p=0.98; R^2^=8.25×10^-6^. We carried out co-variation analyses using package GGally (Figure S3) and fitted a single GLM to the full dataset to avoid multiple testing. As only four (N-1) variables of community member proportion could be fitted to the data simultaneously, we initially constructed five binomial GLMs describing pKJK5-bearing *Variovorax* fraction as a function of Treatment and four community constituent proportions, leaving out each in turn. In these models, *Ochrobactrum* proportion never reached statistical significance.

Therefore, we constructed a GLM lacking this variable as a starting point and successively reduced the model to only include significant variables. See Table S3 for a summary of which additional model variables were dropped due to insignificance. The final model was a binomial GLM with logit link function: Plasmid-bearing *Variovorax* fraction was modelled as a function of Treatment (community), *Stenotrophomonas* proportion, and *Achromobacter* proportion. Total *Variovorax* colony counts for each sample were taken into account as weights. F =0.4635, df=17 & 62, pseudo R^2^=0.848.

#### Plasmid maintenance-fitness correlation (Figure 4)

The dataset combines selection coefficient data (Figure 1) with plasmid maintenance data (Figure 2). To synthesise matching treatments, we built a general linear mixed-effects model using replicates from each experiment as random effects. This was done to statistically investigate all datapoints, rather than just arithmetic means. Only arithmetic means and standard deviation are displayed in Figure 4 because aligning these metrics for each replicate across the two experiments is not meaningful.

pKJK5-bearing *Variovorax* fraction was modelled as a function of plasmid selection coefficient, with random intercept effects of fitness experiment replicate and maintenance experiment replicate. F=5.2581; 25 observations with 2x 5 random-effect groups, conditional R^2^ = marginal R^2^ = 0.1797. Chi-test of inclusion of selection coefficient confirmed this variable is a significant constituent of the model; p = 0.023.

#### Community maintenance experiment (Figure 5)

A single extremely influential datapoint was removed for statistical analyses (P+pKJK5::*gfp* community treatment at T5; the only replicate where *Pseudomonas* was detected with one GFP-colony). Some monoculture samples were contaminated with colonies of other species (2 samples at T5, 4 samples at T17). These replicates were entirely removed for all data visualisation and analyses, resulting in N=4-5 for all treatments.

We constructed a binomial GLM with logit link function for each timepoint: pKJK5-bearing colony fraction was modelled as a function of Species, culture conditions (monoculture / community), plasmid type (pKJK5::*gfp* / pKJK5::*gfp*^PL^), and their interactions. Total focal species colony counts were taken into account for each sample as weights. T5: F=1.487, df=13 & 73, pseudo R^2^=0.996. T17: F=0.6397, df=19 & 74, pseudo R^2^=0.979.

## Abbreviations

Antimicrobial resistance (AMR): resistance of a microbe to an antibiotic or antimicrobial drug
Generalised Linear Model (GLM): statistical model
Linear Model (LM): statistical model
Honest Signficance Difference test (HSD): Tukey’s HSD; testing for significant differences between individual treatments after fitting a statistical model.
Tryptone Soy Broth (TSB): Growth Medium

## Acknowledgments & Funding

We thank Suzanne Kay and Meaghan Castledine for kindly providing *Pseudomonas* sp., *Stenotrophomonas* sp., *Achromobacter* sp., *Ochrobactrum* sp., and *Variovorax* sp. isolates, Arthur Newbury for providing pKJK5*::gfp* PSAOV transconjugants, and Angus Buckling for insights into the community model system. Additionally, we thank Elizabeth Pursey for helpful advice on statistical analyses. We thank Stefano Pagliara and James Hall (University of Liverpool, UK) for insightful discussions around community composition data analysis.

DWS was supported in part by grant MR/N0137941/1 for the GW4 BIOMED MRC DTP, awarded from the Medical Research Council (MRC)/UKRI.

UK was supported by the ANTIVERSA project funded by the German Bundesministerium für Bildung, und Forschung under grant number 01LC1904A. Responsibility for the information and views expressed in the manuscript lies entirely with the authors.

SVH acknowledges funding from the Biotechnology and Biological Sciences Research Council (BB/R010781/1; BB/S017674/1) and the Lister Institute for Preventative Medicine.

## Supporting Information Captions

<u>Supplementary Data:</u> Raw colony counts and selection coefficient data from the fitness experiment (Figure 1), raw colony counts from the community maintenance experiment (Figure 2), and raw colony counts from the final community maintenance experiment (Figure 5). Available in separate datasheets as a supplementary excel file.

<u>Table S1: Conjugation efficiencies to *Variovorax* during the fitness experiment.</u> Conjugation efficiency, defined as the final proportion of transconjugants within the recipient population, after 5 days co-culture is given for each sample. Other species did not form transconjugants.

<u>Table S2: Growth partner experimental values.</u> P value refers to the probability of the treatment average being significantly different from the average of treatment 1.1 as assessed by the statistical model in **Figure 2**.

<u>Table S3: Growth partner experiment statistical model details.</u> This table outlines the constituents of a statistical model fitted to the data of the experiment testing *Variovorax* plasmid maintenance in the presence of various growth partners. Significance values of individual model constituents were derived by Chi test. This model is a binomial model with logit link function and was further reduced by removal of non-significant (Pr>0.05) constituents for final data analysis for Figure 3.

Model function: V_fraction ~ Treatment + replicate + comp_P + comp_S + comp_A + comp_O

As only four (N-1) explanatory variables of species proportion could be fitted to the statistical model simultaneously (N=5), the probability of significance for comp_V was assessed after removal of the variable comp_O. This variable was chosen for removal as it did not reach significance with any combination of included explanatory variables.

<u>Figure S1: pKJK5 variants.</u> Block arrows indicate ORFs, length not to scale. **a WT pKJK5**. Published by [27], genbank acc. AM261282.1. The indicated section is accessory gene load 2 spanning from nts 21682-33379. **b pKJK5::*gfp*^PL^** payload is inserted at position 22540 within *intI1* and consists of SpyCas9 and non-targeting sgRNA with constitutive promoters as well as GFPmut3b as in c. **c *pKJK5::gfp*** payload is inserted at position 23107 and consists of GFPmut3b with lacI-repressible promoter. Published by [6].

<u>Figure S2: Design of Growth Partner Experiment.</u> **A Strains at T0.** *Variovorax* is the only strain which carries pKJK5::*gfp*^PL^. **B Various community treatments.** *Variovorax* was passaged in monoculture (1.1) or with 1-4 growth partners. **C Experimental setup.** T0 strains were co-incubated (in absence of selection) and transferred at T3. At T5, communities were plated onto KB agar and community composition was determined by counting colony morphologies. pKJK5-bearing *Variovorax* fraction was determined by analysing GFP expression on plates using a fluorescence lamp (not shown).

<u>Figure S3: Pairwise correlations of community composition.</u> Proportion of each species within each community plotted against each other; datapoints with community proportions containing 0 in each panel were removed (compare **Figure 3**). On the diagonal, a density line indicates the data point distribution of each species on an arbitrary scale. Above the diagonal, Pearson correlation coefficients are given. Significant pairwise correlations are indicated (*/**/***) and highlighted in boxes. *Variovorax* proportion was found to increase together with the presence of each growth partner; analogous to observed *Variovorax* density increases in pairwise competitions [16]. *Achromobacter* proportion had a negative relationship with *Pseudomonas* and *Stenotrophomonas* proportion, which may in part explain the opposing effects these species have on *Variovorax* plasmid maintenance.

P – *Pseudomonas*; S – *Stenotrophomonas*; A – *Achromobacter*; O – *Ochrobactrum*; V – *Variovorax*.

## Notes

### Competing Interest Statement

The authors have declared no competing interest.

## References

1. Koonin EV. Horizontal gene transfer: essentiality and evolvability in prokaryotes, and roles in evolutionary transitions. F1000Research. 2016;5: 1805. doi:10.12688/f1000research.8737.1

2. Brockhurst MA, Harrison E, Hall JPJ, Richards T, McNally A, MacLean C. The Ecology and Evolution of Pangenomes. Curr Biol. 2019;29: R1094–R1103. doi:10.1016/j.cub.2019.08.012

3. Partridge SR, Kwong SM, Firth N, Jensen SO. Mobile genetic elements associated with antimicrobial resistance. Clin Microbiol Rev. 2018;31. doi:10.1128/CMR.00088-17

4. Elwell LP, Shipley PL. Plasmid-Mediated Factors Associated with Virulence of Bacteria to Animals. Annu Rev Microbiol. 1980;34: 465–496. doi:10.1146/annurev.mi.34.100180.002341

5. Dewar AE, Thomas JL, Scott TW, Wild G, Griffin AS, West SA, et al. Plasmids do not consistently stabilize cooperation across bacteria but may promote broad pathogen host-range. Nat Ecol Evol. 2021;5: 1624–1636. doi:10.1038/s41559-021-01573-2

6. Klümper U, Riber L, Dechesne A, Sannazzarro A, Hansen LH, Sørensen SJ, et al. Broad host range plasmids can invade an unexpectedly diverse fraction of a soil bacterial community. ISME J. 2015;9: 934–945. doi:10.1038/ismej.2014.191

7. San Millan A, MacLean RC. Fitness Costs of Plasmids: a Limit to Plasmid Transmission. Microbiol Spectr. 2017;5: 5.5.02. doi:10.1128/microbiolspec.MTBP-0016-2017

8. Hall JPJ, Harrison E, Lilley AK, Paterson S, Spiers AJ, Brockhurst MA. Environmentally co-occurring mercury resistance plasmids are genetically and phenotypically diverse and confer variable context-dependent fitness effects. Environ Microbiol. 2015;17: 5008–5022. doi:10.1111/1462-2920.12901

9. Alonso-del Valle A, León-Sampedro R, Rodríguez-Beltrán J, DelaFuente J, Hernández-García M, Ruiz-Garbajosa P, et al. Variability of plasmid fitness effects contributes to plasmid persistence in bacterial communities. Nat Commun. 2021;12: 2653. doi:10.1038/s41467-021-22849-y

10. Hall JPJ, Wood AJ, Harrison E, Brockhurst MA. Source–sink plasmid transfer dynamics maintain gene mobility in soil bacterial communities. Proc Natl Acad Sci. 2016;113: 8260-8265.

11. Stewart FM, Levin BR. The Population Biology of Bacterial Plasmids: A PRIORI Conditions for the Existence of Conjugationally Transmitted Factors. Genetics. 1977;87: 209-228.

12. Bergstrom CT, Lipsitch M, Levin BR. Natural selection, infectious transfer and the existence conditions for bacterial plasmids. Genetics. 2000;155: 1505–1519.

13. Alseth EO, Pursey E, Luján AM, McLeod I, Rollie C, Westra ER. Bacterial biodiversity drives the evolution of CRISPR-based phage resistance. Nature. 2019;574: 549–552. doi:10.1038/s41586-019-1662-9

14. Klümper U, Recker M, Zhang L, Yin X, Zhang T, Buckling A, et al. Selection for antimicrobial resistance is reduced when embedded in a natural microbial community. ISME J. 2019;13: 2927–2937. doi:10.1038/s41396-019-0483-z

15. Castledine M, Padfield D, Buckling A. Experimental (co)evolution in a multi-species microbial community results in local maladaptation. Ecol Lett. 2020;23: 1673–1681. doi:10.1111/ele.13599

16. Padfield D, Castledine M, Pennycook J, Hesse E, Buckling A. Short-term relative invader growth rate predicts long-term equilibrium proportion in a stable, coexisting microbial community. bioRxiv [preprint]; 2020 [cited 2022 May 20]. doi:10.1101/2020.04.24.059097. Available from: https://www.biorxiv.org/content/10.1101/2020.04.24.059097v1.full

17. Jiang Y, Qian F, Yang J, Liu Y, Dong F, Xu C, et al. CRISPR-Cpf1 assisted genome editing of Corynebacterium glutamicum. Nat Commun. 2017;8. doi:10.1038/ncomms15179

18. Zhang J, Zong W, Hong W, Zhang Z-T, Wang Y. Exploiting endogenous CRISPR-Cas system for multiplex genome editing in Clostridium tyrobutyricum and engineer the strain for high-level butanol production. Metab Eng. 2018;47: 49–59. doi:10.1016/j.ymben.2018.03.007

19. Cho S, Choe D, Lee E, Kim SC, Palsson B, Cho B-K. High-Level dCas9 Expression Induces Abnormal Cell Morphology in Escherichia coli. ACS Synth Biol. 2018;7: 1085–1094. doi:10.1021/acssynbio.7b00462

20. Bahl MI, Hansen LH, Sørensen SJ. Impact of conjugal transfer on the stability of IncP-1 plasmid pKJK5 in bacterial populations. FEMS Microbiol Lett. 2007;266: 250–256. doi:10.1111/j.1574-6968.2006.00536.x

21. Lopatkin AJ, Meredith HR, Srimani JK, Pfeiffer C, Durrett R, You L. Persistence and reversal of plasmid-mediated antibiotic resistance. Nat Commun. 2017;8: 1689. doi:10.1038/s41467-017-01532-1

22. Kottara A, Hall JPJ, Brockhurst MA. The proficiency of the original host species determines community-level plasmid dynamics. FEMS Microbiol Ecol. 2021;97: fiab026. doi:10.1093/femsec/fiab026

23. Kottara A, Carrilero L, Harrison E, Hall JPJ, Brockhurst MAY 2021. The dilution effect limits plasmid horizontal transmission in multispecies bacterial communities. Microbiology. 2021;167: 001086. doi:10.1099/mic.0.001086

24. Zhang W, Bielaszewska M, Kunsmann L, Mellmann A, Bauwens A, Köck R, et al. Lability of the pAA Virulence Plasmid in Escherichia coli O104:H4: Implications for Virulence in Humans. PLOS ONE. 2013;8. doi:10.1371/journal.pone.0066717

25. Li L, Dechesne A, Madsen JS, Nesme J, Sørensen SJ, Smets BF, et al. Plasmids persist in a microbial community by providing fitness benefit to multiple phylotypes. ISME J. 2020;14: 1170–1181. doi:10.1038/s41396-020-0596-4

26. Pursey E, Sünderhauf D, Gaze WH, Westra ER, van Houte S. CRISPR-Cas antimicrobials: Challenges and future prospects. Hogan DA, editor. PLOS Pathog. 2018;14: e1006990. doi:10.1371/journal.ppat.1006990

27. Bahl MI, Hansen LH, Goesmann A, Sørensen SJ. The multiple antibiotic resistance IncP-1 plasmid pKJK5 isolated from a soil environment is phylogenetically divergent from members of the previously established α, β and δ sub-groups. Plasmid. 2007;58: 31–43. doi:10.1016/j.plasmid.2006.11.007

28. Lee DJ, Bingle LE, Heurlier K, Pallen MJ, Penn CW, Busby SJ, et al. Gene doctoring: A method for recombineering in laboratory and pathogenic Escherichia coli strains. BMC Microbiol. 2009;9. doi:10.1186/1471-2180-9-252

29. Jinek M, Chylinski K, Fonfara I, Hauer M, Doudna JA, Charpentier E. A Programmable Dual-RNA-Guided DNA Endonuclease in Adaptive Bacterial Immunity. Science. 2012;337: 816–821. doi:10.1126/science.1225829

30. Ferrières L, Hémery G, Nham T, Guérout AM, Mazel D, Beloin C, et al. Silent mischief: Bacteriophage Mu insertions contaminate products of Escherichia coli random mutagenesis performed using suicidal transposon delivery plasmids mobilized by broad-host-range RP4 conjugative machinery. J Bacteriol. 2010;192: 6418–6427. doi:10.1128/JB.00621-10

31. Martínez-García E, Calles B, Arévalo-Rodríguez M, de Lorenzo V. pBAM1: an all-synthetic genetic tool for analysis and construction of complex bacterial phenotypes. BMC Microbiol. 2011;11: 38. doi:10.1186/1471-2180-11-38

32. Dimitriu T, Kurilovich E, Łapińska U, Severinov K, Pagliara S, Szczelkun MD, et al. Bacteriostatic antibiotics promote CRISPR-Cas adaptive immunity by enabling increased spacer acquisition. Cell Host Microbe. 2022;30: 31–40.e5. doi:10.1016/j.chom.2021.11.014

